# Transhydrogenase and growth substrate influence lipid hydrogen isotope ratios in *Desulfovibrio alaskensis* G20

**DOI:** 10.1101/039792

**Authors:** William D. Leavitt, Theodore M. Flynn, Melanie K. Suess, Alexander S. Bradley

## Abstract

Microbial fatty acids preserve metabolic and environmental information in their hydrogen isotope ratios (^2^H/^1^H). This ratio is influenced by parameters that include the ^2^H/^1^H of water in the microbial growth environment, and biosynthetic fractionations between water and lipid. In some microbes, this biosynthetic fractionation has been shown to vary systematically with central energy metabolism, and controls on fatty acid ^2^H/^1^H may be linked to the intracellular production of NADPH. We examined the apparent fractionation between media water and the fatty acids produced by *Desulfovibrio alaskensis* G20. Growth was in batch culture with malate as an electron donor for sulfate respiration, and with pyruvate and fumarate as substrates for fermentation and for sulfate respiration. A larger fractionation was observed as a consequence of respiratory or fermentative growth on pyruvate than growth on fumarate or malate. This difference correlates with opposite apparent flows of electrons through the electron bifurcating/confurcating transhydrogenase NfnAB. When grown on malate or fumarate, mutant strains of *D. alaskensis* G20 containing transposon disruptions in a copy of *nfnAB* show different fractionations than the wild type strain. This phenotype is muted during fermentative growth on pyruvate, and it is absent when pyruvate is a substrate for sulfate reduction. All strains and conditions produced similar fatty acid profiles, and the ^2^H/^1^H of individual lipids changed in concert with the mass-weighted average. Unsaturated fatty acids were generally depleted in ^2^H relative to their saturated homologues, and anteiso-branched fatty acids were generally depleted in ^2^H relative to straight-chain fatty acids. Fractionation correlated with growth rate, a pattern that has also been observed in the fractionation of sulfur isotopes during dissimilatory sulfate reduction by sulfate reducing bacteria.

**Abbreviations**
CNIH: Cornichon

## 1 Introduction

The structures and isotopic compositions of lipids preserve information about organisms that can be archived in sediments and rocks over geological time scales. Understanding how to interpret this information is a central task of organic geochemistry. Lipid structures can be affiliated to particular groups of organisms (Pearson, 2014), and ratios of carbon isotopes in lipids record information about carbon sources and assimilation pathways (Hayes, 2001; Smith and Epstein, 1970) The ratio of hydrogen isotopes (deuterium/protium = ^2^H/^1^H), in lipids derived from environmental samples has been observed to relate to the ^2^H/^1^H of environmental water (Hayes, 2001; Sauer et al., 2001). More recently, it was shown in a range of aerobic microorganisms that the fractionation of hydrogen isotopes between media water and lipids varied with changes in growth substrate (Zhang et al., 2009). Experiments with anaerobes have shown less systematic changes in lipid ^2^H/^1^H as a function of energy metabolism, although strong differences have been observed in lipid ^2^H/^1^H of organisms in pure culture versus co-cultures with another organism (Dawson et al., 2015; Osburn, 2013; Osburn et al., 2016).

Observations that lipid ^2^H/^1^H varies as a function of growth substrate in many microorganisms raises the question of what specific metabolic mechanisms are responsible. Zhang et al. (2009) considered several explanations, and through a process of elimination, deduced that observed differences in lipid ^2^H/^1^H must be a consequence of differences in the NAD(P)H that serves as a hydride donor during lipid biosynthesis. These authors pointed out that cells have multiple pathways for producing NAD(P)H, and that the relative importance of each of these mechanisms varies with differences in growth conditions. One mechanism considered was alteration of the ^2^H/^1^H ratio of the transferrable hydride in NAD(P)H by transhydrogenase enzymes. This suggestion stems from two key observations. First, up to half of the hydrogen atoms in microbial lipids are derived directly from NADPH during biosynthesis (Jackson, 2003; Saito et al., 1980). Second, in vitro observations of the hydrogen isotope fractionation imparted by transhydrogenase suggest that it is very large (> 800‰) (Bizouarn et al., 1995; Jackson et al., 1999; Venning et al., 1998).

In this study, we vary substrates and use mutant strains to investigate the importance of a transhydrogenase (NADH-dependent reduced ferredoxin:NADP oxidoreductase; NfnAB) on the lipid ^2^H/^1^H ratios in an anaerobic microorganism, *Desulfovibrio alaskensis* G20. Recent work has suggested that NfnAB plays an important role in energy conservation in this microbe (Price et al., 2014). The role of NfnAB varies as a function of the growth substrate. During growth on malate, for example, NfnAB is predicted to catalyze an electron bifurcation reaction (Buckel and Thauer, 2013) in which NADPH reduces ferredoxin and NAD^+^ to produce NADH (Figure 1). Conversely, during growth on pyruvate, NfnAB is predicted to catalyze electron confurcation and the production of NADPH from NADH, NADP^+^, and reduced ferredoxin. The importance of this transhydrogenase to anaerobic energy metabolism may be critical to understanding lipid H-isotope signatures. Since the transhydrogenase reaction catalyzed by NfnAB is predicted to occur in opposite directions during growth on pyruvate versus that on malate, the lipids produced under each condition should have different ^2^H/^1^H ratios if NfnAB is indeed a significant source of isotope fractionation for intracellular hydrogen. Furthermore, the role of NfnAB in hydrogen isotope fractionation can be further explored using mutant strains of *D. alaskensis* G20 in which the NfnAB-2 loci have been disrupted.

**Figure 1.**
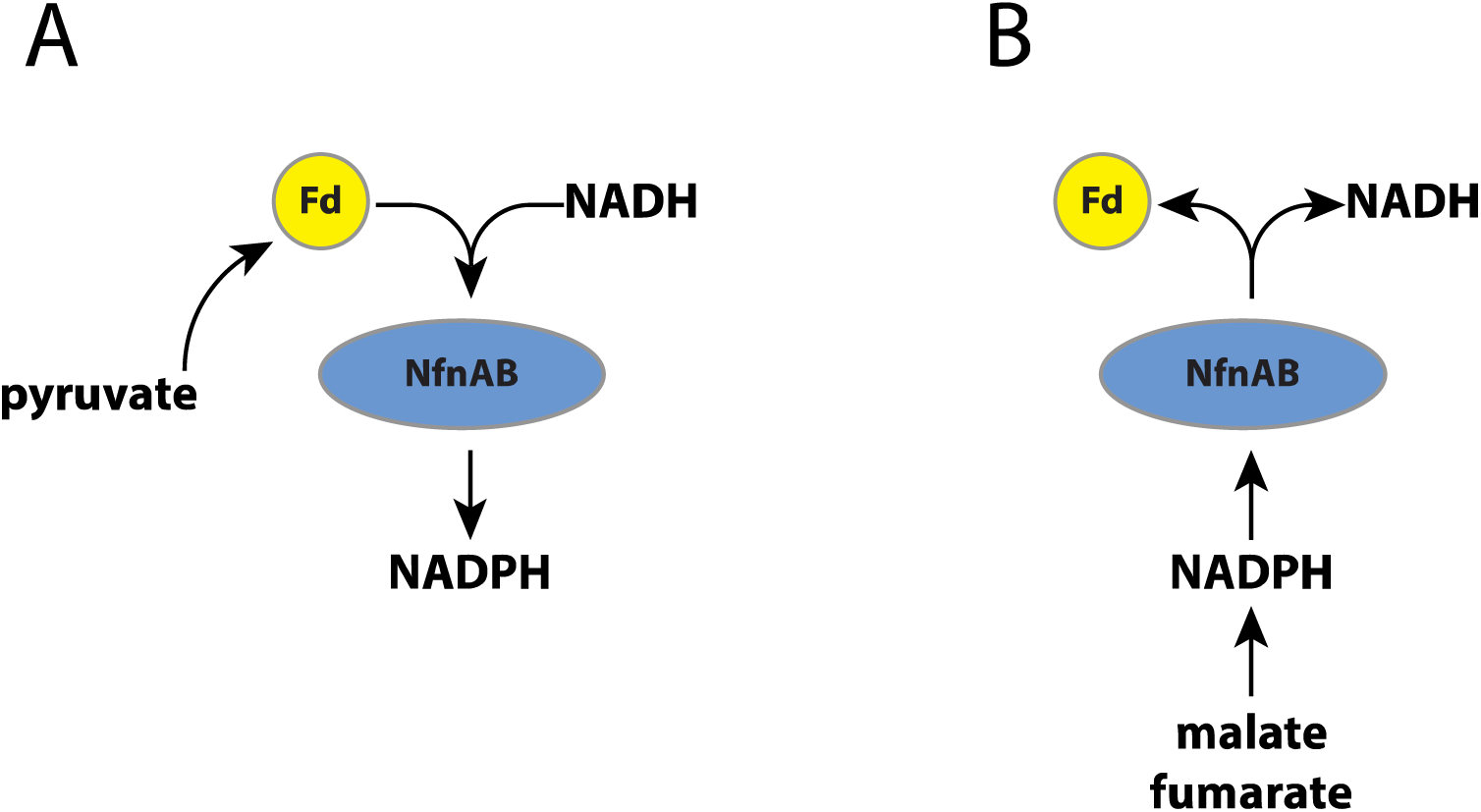
(A). A schematic pathway of electron flow during pyruvate respiration or fermentation in WT *D. alaskensis* G20. Electrons are transferred from pyruvate through ferredoxin and NADPH, and NfnAB catalyzes electron confurcation, producing NADPH. Mutation of NfnAB inhibits this reaction and may result in a relative NADPH deficiency (B). Electron flow during fumarate and malate respiration or fumarate fermentation. Mutation of NfnAB inhibits this reaction and may result in a relative NADPH surplus.

## 2 Material and Methods

### 2.1 Strains, growth media, culture conditions and biomass sampling

Wild type *Desulfovibrio alaskensis* G20 was obtained along with two mutant strains from the library collection at Lawrence Berkeley National Laboratory. Each mutant contains a Tn5 transposon insertion into a gene of interest. These insertions (Kuehl et al., 2014) were into the genes *nfnA*-2 at locus Dde_1250 (strain JK00256) and *nfn*B-2 at locus Dde_1251 (JK01775). Hereafter, these strains are referred to as the *nfnA*-2 and *nfnB*-2 mutants. These loci encode the subunits for one of two paralogs of NfnAB in *D. alaskensis* G20.

All strains were resuscitated from 10% glycerol freezer stocks stored at −80 °C. Resuscitated strains were inoculated into serum bottles containing approximately 50 ml of a rich lactate/sulfate medium containing yeast extract (MOY_LS) and incubated at 30 °C. After reaching stationary phase, strains were then serially transferred three times in a defined lactate/sulfate (80 mM/40 mM) medium (MO_LS). Late-log phase cultures of the third transfer were diluted 1 to 100 into duplicate bottles for each isotope fractionation experiment. There were five experimental growth conditions, which combined an electron donor and 40 mM sulfate (for sulfate respiration), or an electron donor alone for fermentation. The five conditions were: pyruvate/sulfate respiration, malate/sulfate respiration, fumarate/sulfate respiration, pyruvate fermentation, fumarate fermentation.

The basal growth medium recipe (MO) was as follows: 8 mM magnesium chloride, 20 mM ammonium chloride, 0.6 mM calcium chloride, 6 mL/L trace elements solution (see below), 0.12 mM of FeCl_2_ (125mM)+EDTA (250 mM) stock solution, 30 mM Tris-HCl (2M, pH 7.4 stock). Sodium thioglycolate (0.12 g/L) was added as a reductant following initial degassing. MO medium containing yeast extract (MOY) is generated by adding yeast extract (0.1% w/v) to MO medium from an anoxic sterile stock. Media were made anaerobic by degassing with O_2_-free N_2_ that had been filtered through sterile 0.22 μm syringe filters. Solutions were degassed for 2 hours per liter. The pH of the final medium was adjusted to 7.2 using sterile and anoxic HCl or NaOH, autoclave-sterilized, and then cooled under sterile O_2_-free N_2_. After cooling, phosphate solution was added to a final concentration of 2 mM from a sterile, anoxic stock solution of K_2_HPO_4_+NaH_2_PO_4_. Thauer’s Vitamins were added from a 1000x stock (Rabus et al., 2015). The initial concentration of sulfate was always 40 mM (except in fermentation experiments) and was added directly to the medium from a sterile and anoxic stock solution of Na_2_SO_4_ solution. Electron donors (sodium lactate, sodium pyruvate, malic acid, or sodium fumarate) were prepared separately as 1M stocks in MilliQ water, adjusted to a pH of 7.2, and degassed in a manner similar to the basal media. These anoxic stocks were then added to the basal media using aseptic technique.

Growth rate was determined by monitoring changes in optical density (OD_600_) over time for each experiment. Replicate cultures were tracked through log-phase and into early stationary phase, at which point they were harvested for biomass. Growth rates were calculated using a modified logistic equation (Rabus et al., 2006) and averaged across the apparent log-phase of growth. In experiments showing diauxic growth, we calculated a weighted average growth rate, where weighting accounts for the amount of biomass produced during each growth interval.

Duplicate 50mL cultures were harvested at the onset of early stationary phase by opening the serum bottles, decanting the remainder of each serum bottle (> 40mL) into sterile 50mL conical tubes, and centrifuging at 5000 rpm at 5 °C for 30 minutes. Spent medium was decanted into a fresh 50mL tube and frozen at −80 °C for later analysis of the isotopic composition of water therein. The biomass pellet was frozen at −80 °C, transferred to a pre-baked and weighed 4mL borosilicate glass vial, lyophilized, and weighed.

### 2.2 Fatty acid extraction, identification, and quantitation

Samples were simultaneously extracted and derivatized to fatty acid methyl ethers (FAMEs) by adding a mixture of hexane, methanol and acetyl chloride to the lyophilized cell pellet, followed by heating at 100 °C for 10 minutes, and extraction with hexane (Rodriguez-Ruiz et al., 1998; Zhang et al., 2009). This procedure was also concurrently performed on two isotope standards, myristic acid and phthalic acid, for which the δ^2^H of non-exchangeable hydrogen has been determined (Qi and Coplen, 2011). Each sample was reacted with acid-activated copper shot to remove elemental sulfur, and then concentrated under a stream of dry hydrocarbon-free nitrogen.

Individual FAMEs were analyzed using a HP 7890 gas chromatograph fitted with a split/splitless injector operated in splitless mode, equipped with a J&W DB-5 fused silica capillary column (30 m length, 0.25-mm inner diameter, and 0.25-μm film thickness) and coupled to an Agilent 6973 mass selective detector. FAME identifications were based on mass spectra and retention times and are reported in Table S1. Retention times were converted to Kovats retention indices by comparison to a mix of *n*-alkanes, and compared to the retention indices of published fatty acids (Dickschat et al., 2011; Taylor and Parkes, 1983). Double bond locations were identified by converting unsaturated fatty acids to their dimethyl disulfide adducts (Nichols et al., 1986). Abundances were determined by peak area as calculated in Chemstation (Agilent Technologies, Santa Clara, CA) relative to a known amount of co-injected methyl tetracosanoate (C24:0).

### 2.3 Isotopic measurements and data handling

Hydrogen-isotopic compositions of individual FAMEs were determined using a TraceGC gas chromatograph fitted with a column identical to that on the Agilent GC, and coupled to a Thermo Scientific Delta V Plus isotope-ratio-monitoring mass spectrometer via a Thermo GC-Isolink pyrolysis interface at 1400 °C. Column temperature was initially 60 °C and was increased at a rate of 6 °C min^−1^ until reaching a final temperature of 320 °C. Hydrogen isotope ratios of individual lipids were determined relative to coinjected methyl tetracosanoate (C24:0) of known isotopic composition, provided by Dr. A. Schimmelmann (Indiana University). Instrumental precision was regularly monitored by analyzing the δ^2^H on external standard mixtures of FAMEs and of *n*-alkanes with previously determined isotopic composition (V-SMOW), also purchased from Dr. Schimmelmann (Indiana University). Over the measurement period the mean RMS error on a mixture of 8 FAMEs was 5.5‰ (n = 286). Samples were discarded if they were not bracketed by injections of FAMEs mixture with an RMS better than 7‰. H_3_ factors were determined daily and had a mean value of 2.98 ± 0.3 ppm/nA. All FAME isotopic compositions were corrected by mass balance for the hydrogen present in the methyl group, calculated from them myristic acid and phthalic acid isotopic standards. Samples were reinjected (pseudoreplicates) three to six times, and errors were propagated following established methods (Polissar and D’Andrea, 2014). Statistical analyses were performed in either Prism (GraphPad Software, Inc., La Jolla, CA) or R (RCoreTeam, 2015).

All ^2^H/^1^H ratios are reported as δ^2^H values relative to V-SMOW, and fractionations are reported as apparent fractionations between media water and lipid by the equation: ^2^ε_lipid-water_ = (α_lipid-water_ - 1), where α = [(δ ^2^H_lipid_ + 1)/(δ^2^H_water_ + 1)] and are reported in ‰. The _δ_^2^H_water_ of growth media water was measured using a Picarro L2130-*i* cavity ring-down spectrometer at Northwestern University.

### 2.4 Comparative analysis of *nfnAB* sequences

We constructed a gene tree of *nfnAB* sequences by retrieving data from two public repositories of annotated genomes: the SEED (Overbeek et al., 2014) and IMG (Markowitz et al., 2012). For this study, we restricted sequences to those also found in the subsystem “NADH-dependent reduced ferredoxin:NADP+ oxidoreductase” in the SEED (197 total sequences) and manually refined the list to sequences from known sulfate reducers, methanogens, and other anaerobes. Sequences from organisms known to contain *nfnAB* but not present in the SEED were added manually using a targeted amino acid BLAST search of that organism’s genome in the IMG database. This resulted in a total of 105 *nfnAB* sequences closely related to that of *D. alaskensis* G20 using established criteria (Marti-Renom et al., 2000; Rost, 1999), with > 40% amino acid identity and BLASTP percent identities ranging from 45–77%. *D. alaskensis* G20 itself has two copies of *nfnAB* that share 85% amino acid identity. A multiple sequence alignment of the 105 putative *nfnAB* sequences was created using MUSCLE (Edgar, 2004) and checked manually using AliView (Larsson, 2014). Alignments in FASTA format are available to download in the Supplementary Materials. Pairwise distances for construction of a phylogenetic tree were calculated using the RAxML maximum likelihood algorithm (Stamatakis, 2006) with the program raxmlGUI (Silvestro and Michalak, 2012). The tree itself was generated using the Interactive Tree of Life software (Letunic and Bork, 2011).

## 3. Results

### 3.1 Growth rates

Growth experiments revealed distinct physiological and isotopic phenotypes among the wild type and mutant strains of *D. alaskensis* G20. Growth rates are reported in Table 1 and growth curves are plotted in Figure 2.

Some growth conditions showed clear phenotypic differences between the *D. alaskensis* G20 wild type and the two *nfnAB-2* mutants. Each strain was able to grow as a sulfate reducer using malate as an electron donor, but the growth rate of the mutants was only 22% that of the wild type. Similarly, with fumarate as an electron donor coupled to sulfate reduction, mutant growth rate was roughly 10% that of the wild type. A repression in growth rate (22% of wild type) was also apparent when the strains were grown as fumarate fermenters, in the absence of sulfate. Under all of these conditions, the final optical density of the mutant cultures was less than that of the wild type (Figure 2). The mutant strains exhibited diauxic growth under each of these growth conditions whereas the wild type did not (Figure 2).

**Table 1:** Interval-weighted growth rates of *D. alaskensis* G20 wild type and *nfnAB-2* transhydrogenase mutants on different substrates during sulfate respiration or substrate fermentation. The range in rate is larger for experiments that exhibited bi-phasic (diauxic) growth patters (e.g. fumarate + sulfate), as is apparent from the growth curves (Fig. 2).

| Average growth rates |  |  |
| --- | --- | --- |
| Treatment | Strain | $\mu_{avg} \pm 1\sigma$<br>(per hour) |
| pyruvate +<br>sulfate | wildtype | $0.154 \pm 0.029$ |
| | *nfnA-2* | $0.172 \pm 0.029$ |
| | *nfnB-2* | $0.168 \pm 0.035$ |
| malate +<br>sulfate | wildtype | $0.067 \pm 0.007$ |
| | *nfnA-2* | $0.014 \pm 0.006$ |
| | *nfnB-2* | $0.015 \pm 0.004$ |
| fumarate +<br>sulfate | wildtype | $0.085 \pm 0.012$ |
| | *nfnA-2* | $0.011 \pm 0.002$ |
| | *nfnB-2* | $0.007 \pm 0.006$ |
| fumarate<br>fermentation | wildtype | $0.057 \pm 0.007$ |
| | *nfnA-2* | $0.010 \pm 0.002$ |
| | *nfnB-2* | $0.015 \pm 0.004$ |
| pyruvate<br>fermentation | wildtype | $0.131 \pm 0.007$ |
| | *nfnA-2* | $0.120 \pm 0.001$ |
| | *nfnB-2* | $0.149 \pm 0.012$ |

**Figure 2.**
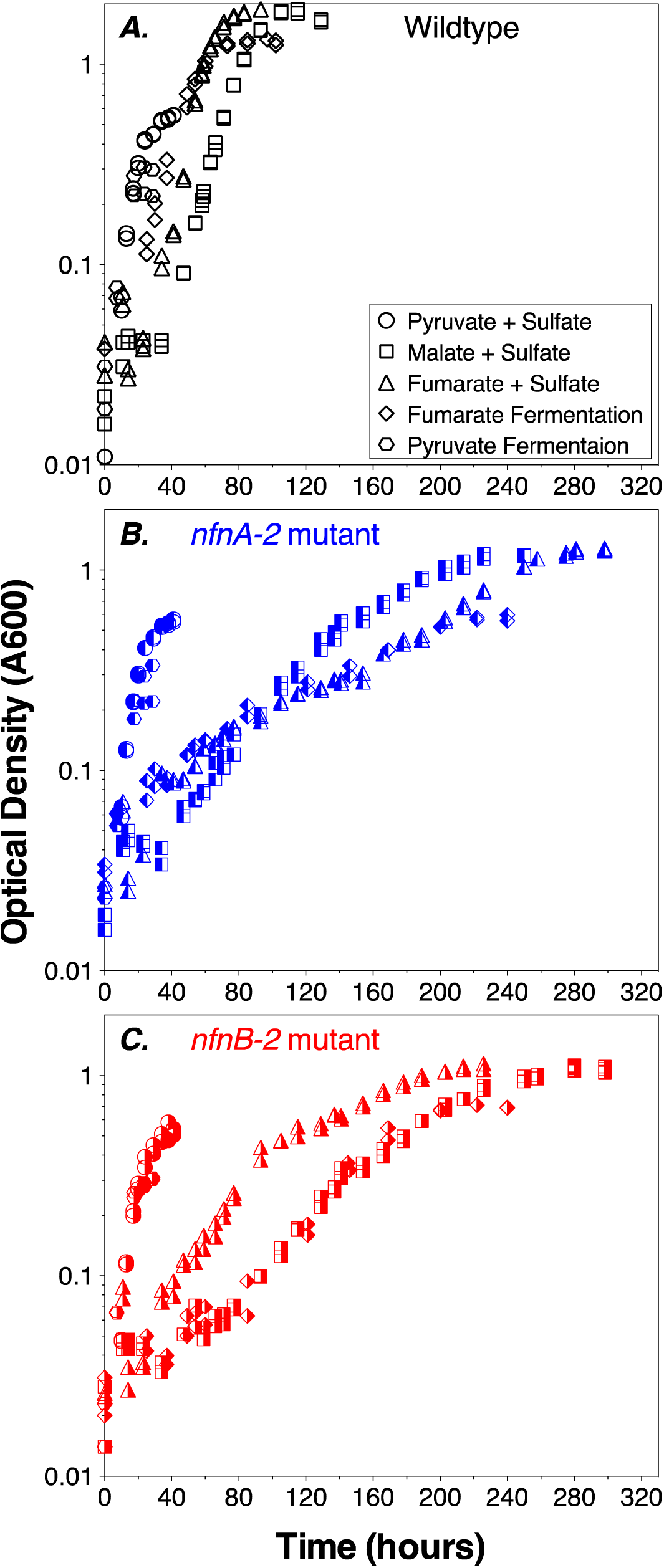
Growth curves for *D. alaskensis* G20 wildtype and transhydrogenase mutants. (A) Wildtype, (B) *nfnA-2* transposon mutant, and (C) *nfnB*-2 transposon mutant, each under all 5-condition sets tested (denoted in the legends). Each symbol represents the average of biological duplicate 50mL cultures. Error bars are smaller than the symbols in all cases. Samples for lipid and isotopic analysis were extracted after the final time-point indicated on this plot, all in early stationary phase.

### 3.2 Lipid profiles

We quantified the abundance of fatty acid structures in each of the three strains under all five experimental conditions (pyruvate/sulfate, malate/sulfate, fumarate/sulfate, fumarate fermentation, or pyruvate fermentation). Fatty acids ranged in carbon number from 14 to 18, and both saturated and monounsaturated fatty acids were present. Branched-chain fatty acids of the iso‐ and anteiso‐ series are present in all three strains under all five experimental conditions. Branched fatty acids contained a total of 15 to 18 carbons, and in some cases contained a double bound.

Differences in the lipid profiles of the mutant relative to the wild type were apparent only under the three conditions in which the mutant showed a growth defect: malate/sulfate, fumarate/sulfate, and fumarate fermentation (Figures 3b, 3c, 3d). During growth on malate/sulfate, the *nfn*AB-2 mutant strains contain a higher proportion of anteiso-C17:0, and a lower proportion of C16:0, C18:0, and iso-C15:0 fatty acids. A similar pattern is seen in the mutants during growth on fumarate/sulfate and during fumarate fermentation, although the C18 patterns are slightly different. Differences in the abundance of anteiso-C17:0 fatty acid is most pronounced in these three growth conditions. In contrast, during both respiratory and fermentative growth on pyruvate, the fatty acid profile of the wild type and mutants were nearly identical (Figures 3a, 3e). Across all strains and conditions, there is a weak inverse correlation between the proportion of branched fatty acids (Figure S1a) or the ratio of anteiso‐ to iso‐ branched compounds (Figure S1b), in each case relative to the mass weighted fractionation. Data used to generate these plots are deposited in a permanent repository at Figshare: doi:10.6084/m9.figshare.2132731.

**Figure 3.**
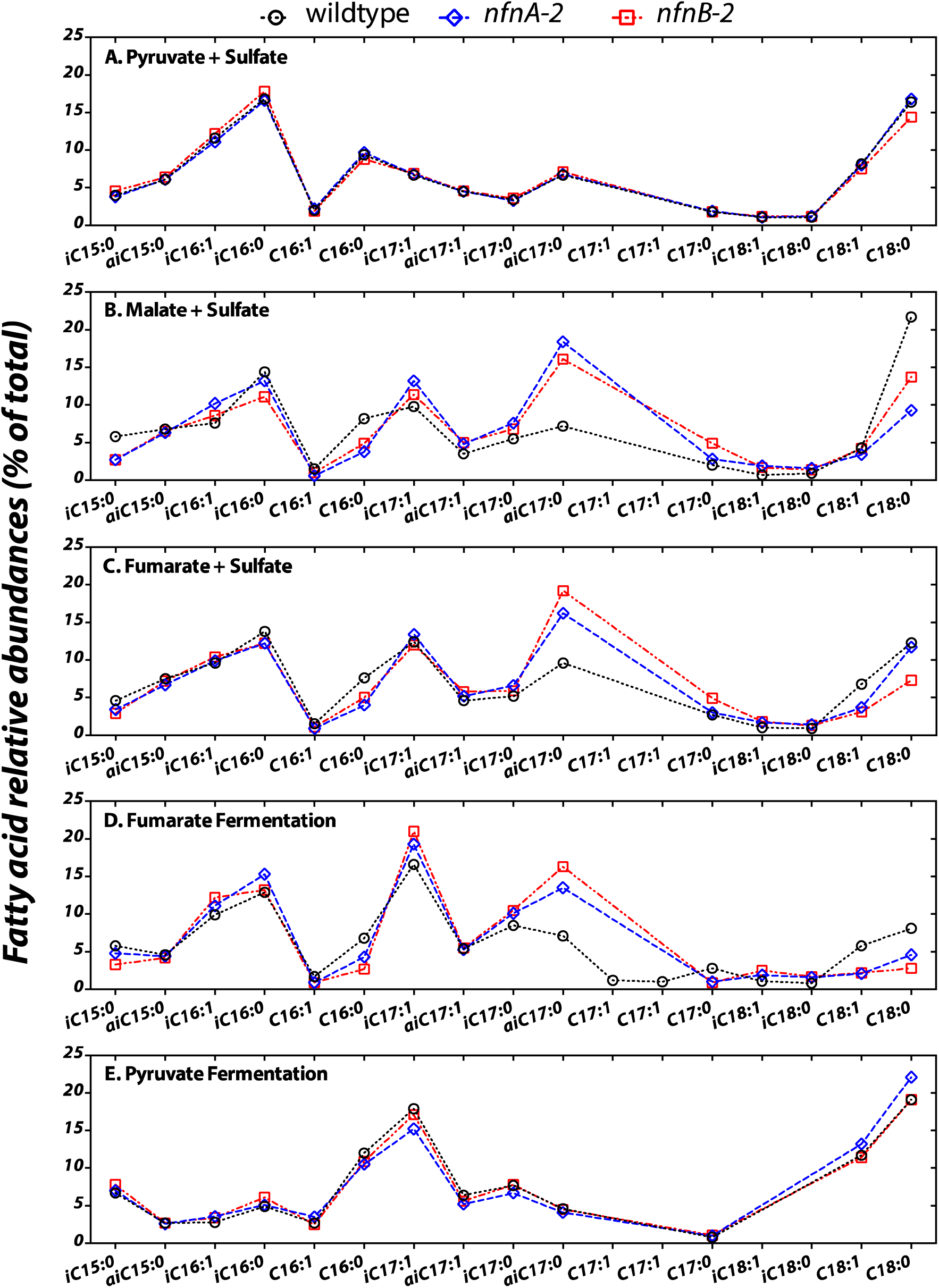
The relative abundance of each fatty acid (% of total) from each strain under the five different growth condition tested (A to E). Sample key: wild type (black circles, dotted lines) or *nfnAB-2* transhydrogenase insertion-deletion mutants, *nfnA-2* (blue diamonds, dashed lines) and *nfnB-2* (red squares, dash-dotted lines). The double bond in *n-*C16, *iso-*C16, *iso-*C17, *anteiso*-C17 is located at the Δ9 carbon, and on the *n*-C18 and *iso*-C18 at the Δ11 carbon. The two C17:1’s were in too low abundance following the DMDS reaction to determine bond positions. Each symbol is the average of biological replicates for each strain given that condition set, and the standard error of individual fatty acid quantifications is <0.5% between biological replicates (error bar significantly smaller than the symbols).

### 3.3 Lipid ^2^H/^1^H fractionations

We calculated δ^2^H_total_ as the weighted average of the δ^2^H_lipid_ of each individual fatty acid pool produced in each strain. We then calculated a total apparent fractionation (^2ε^_total_) for the fatty acid pool (Sessions and Hayes, 2005). The results are shown in Figure 4.

**Figure 4.**
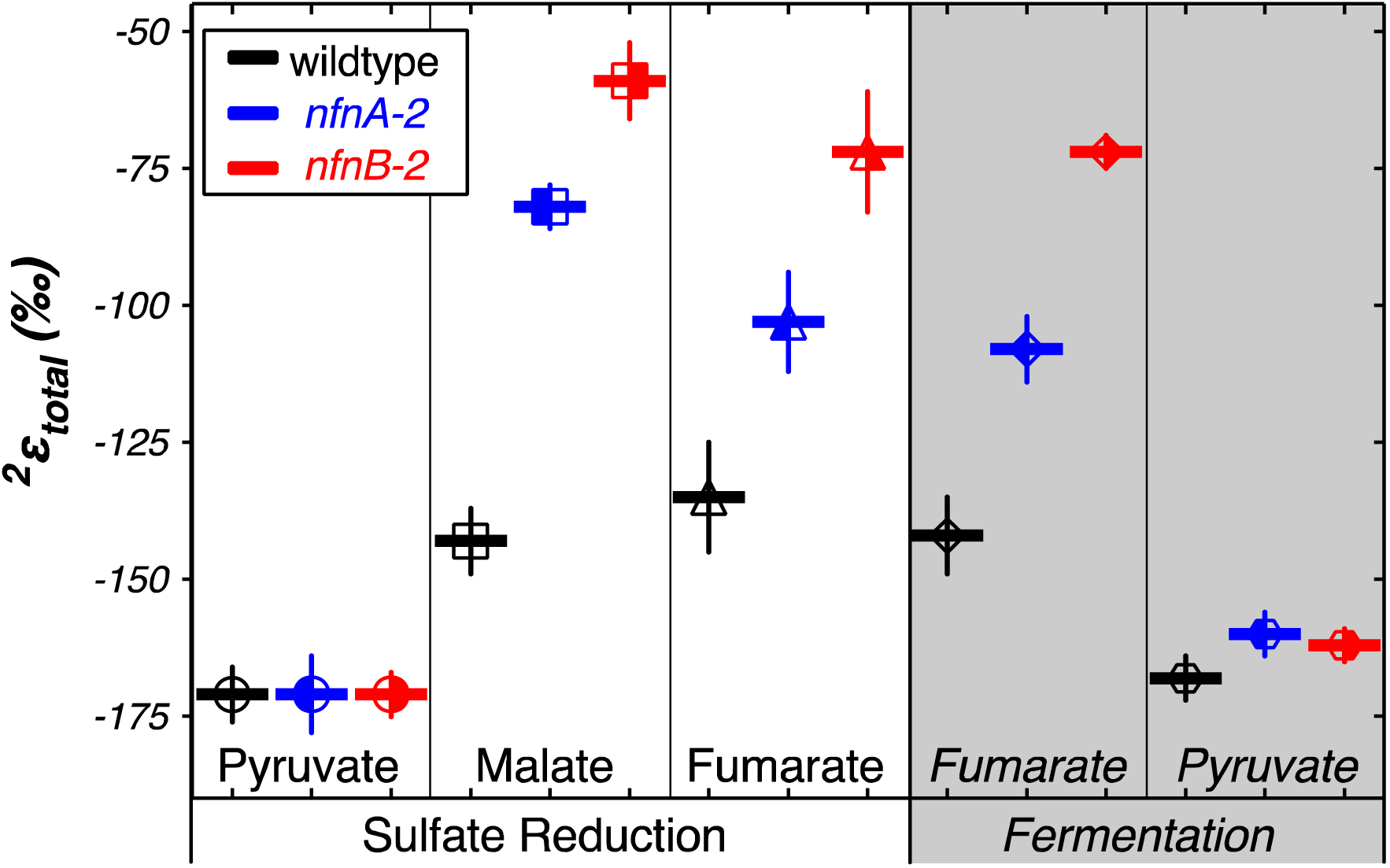
Hydrogen isotope fractionation between media water (symbols) and the mass-weighted lipid pool (^2^ε_total_) for each treatment (horizontal bars). Vertical bars are standard errors on the mean (SEM).

Apparent fractionations produced by the wild type strain were not discernable between pyruvate/sulfate respiration (^2^ε_total_ = −171‰) and pyruvate fermentation (^2^ε_total_ = −168‰). Similarly, both *nfn* mutants have ^2^ε_total_ = −171‰ when grown by pyruvate/sulfate respiration. However, *nfn* mutants that grew by fermenting pyruvate had ^2ε^ for the *nfnA*-2 mutant and ^2^ε_total_ = −162‰ for the *nfnB*-2 mutant.

Differences in ^2^ε_total_ were more pronounced in the other growth conditions. In comparison to growth on pyruvate, the wild type strain showed less negative ^2^ε_total_ as a consequence of malate/sulfate growth (^2^ε_total_ = −143‰), fumarate/sulfate growth (^2^ε_total_ = −135‰), and fumarate fermentation (^2^ε_total_ = −142‰). The *nfn* mutants showed even stronger isotopic phenotypes. The *nfnA*-2 mutant had less negative ^2^ε_total_ than the wild type during growth on malate/sulfate (^2^ε_total_ = −82‰), fumarate/sulfate (^2^ε_total_ = −103‰), and fumarate (^2^ε_total_ = −108‰). The magnitude of fractionation by the *nfnB*-2 mutant was consistently less than that of both the wild type and the *nfnA*-2 mutant on malate/sulfate (^2^ε_total_ = −59‰), fumarate/sulfate (^2^ε_total_ = −72‰), and fumarate (^2^ε_total_ = −72‰).

The δ^2^H_lipid_ of individual lipids can help explain some of these patterns. Most lipids from our cultures were depleted in deuterium by between −50‰ and −250‰ relative to the water in the growth medium. Figures 5 summarize the results from each strain. The various lipid structures produced by each strain had a wide range of ^2^ε_lipid_, but isotopic ordering among lipids was remarkably consistent. Figure 5 shows ^2^ε_lipid_ values for the most abundant lipids in each combination of strain and culture conditions. For all three strains, across nearly every culture condition, the fatty acid with the largest ^2^ε_lipid_ was C16:1. The exceptions to this were produced by the pyruvate fermentation experiments, in which the largest ^2^ε_lipid_ observed was in anteiso-C17:1 in all three strains, and by fumarate fermentation by the mutants, where the C16:1 was in too low abundance for δ^2^H measurements (Figure 5). Across all sulfate reduction experiments, fatty acids containing a double bond were nearly always depleted relative to their saturated homologue. This was true for both straight-chain and branched fatty acids. Differences in δ^2^H were larger between C16 and C16:1 than between C18 and C18:1, driven by the particularly strong ^2^H depletion in C16:1. This pattern was muted in the fermentation experiments. We also found that saturated anteiso-branched fatty acids produced during sulfate reduction were always depleted relative to straight-chain fatty acids. This pattern did not hold for unsaturated anteiso-fatty acids or for lipids produced during fermentation. Saturated iso-branched fatty acids had similar δ^2^H as saturated straight-chain fatty acids. The lipid with the least negative ^2^ε_lipid_ was iso-C18:0 (Figure 5). When produced under certain sulfate reducing conditions this lipid was enriched in ^2^H relative to media water (^2^ε_lipid_ > 0‰). Under fermentative conditions, this lipid was produced in insufficient abundance to measure δ^2^H_lipid_.

**Figure 5.**
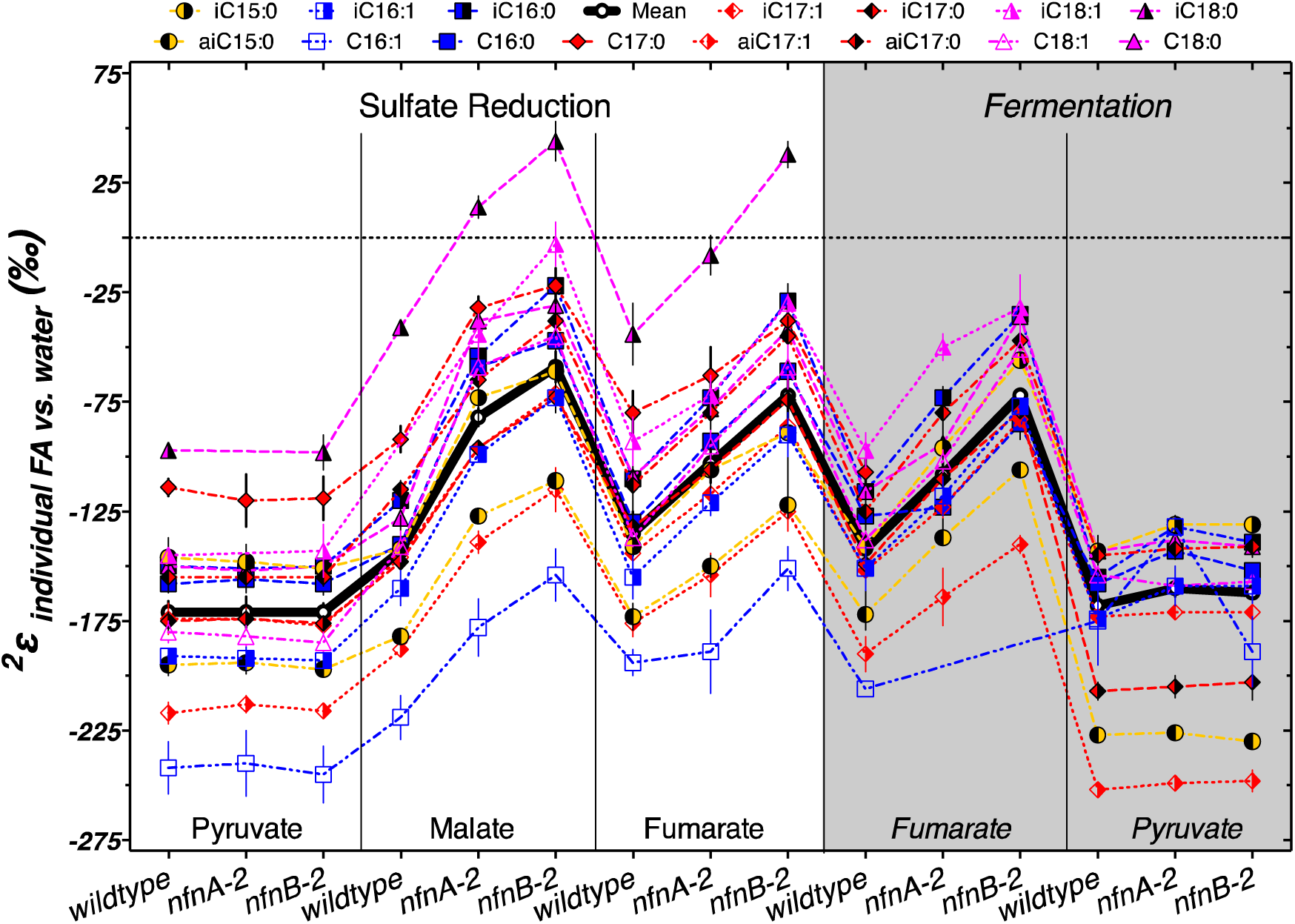
Hydrogen isotope values for individual fatty acids relative to the medium water (symbols) and the mass-weighted lipid pool (^2^ε_total_) for each treatment (horizontal black bars). Each symbol represents the mean of biological replicates (N = 2, and technical replication *n*_avg._ = 3, range 1 to 6), with SEM.

The regular ordering of lipid δ^2^H values suggests that the variations in ^2^ε_total_ were mainly a function of a systematic change in ^2^ε from one condition to another rather than changes in the relative proportion of individual lipids that are particularly enriched or depleted. In Figure 5 the thick black line and circles show ^2^ε_total_, highlighting the relationship between ^2^ε_lipid_ and ^2^ε_total_ for each strain and culture condition. With few exceptions, a consistent pattern emerged in the relative fractionation of each lipid relative to the weighted average. Together, Figures 4 and 5 indicate the presence of significant differences between the wildtype and mutants for growth on malate/sulfate, fumarate/sulfate or fumarate fermentation, while little to no difference existed between strains grown on pyruvate/sulfate or pyruvate fermentation.

We examined whether changes in the abundance of particular lipids were correlated with each other, with growth rate, or with ^2^ε_total_. A graphical display of Pearson correlation indices for each variable pair is shown in Figure S3 This indicates that ^2^ε_total_ is strongly correlated with the relative abundance anteiso-C17:0 fatty acid, and negatively correlated with C16 and C16:1 fatty acid. However, the relative abundance of each of these fatty acids was strongly correlated (negatively, for anteiso-C17:0 fatty acid) with average growth rate (μ). Growth rate emerged as a strong correlate of ^2^ε_total_ (Figure 6).

**Figure 6.**
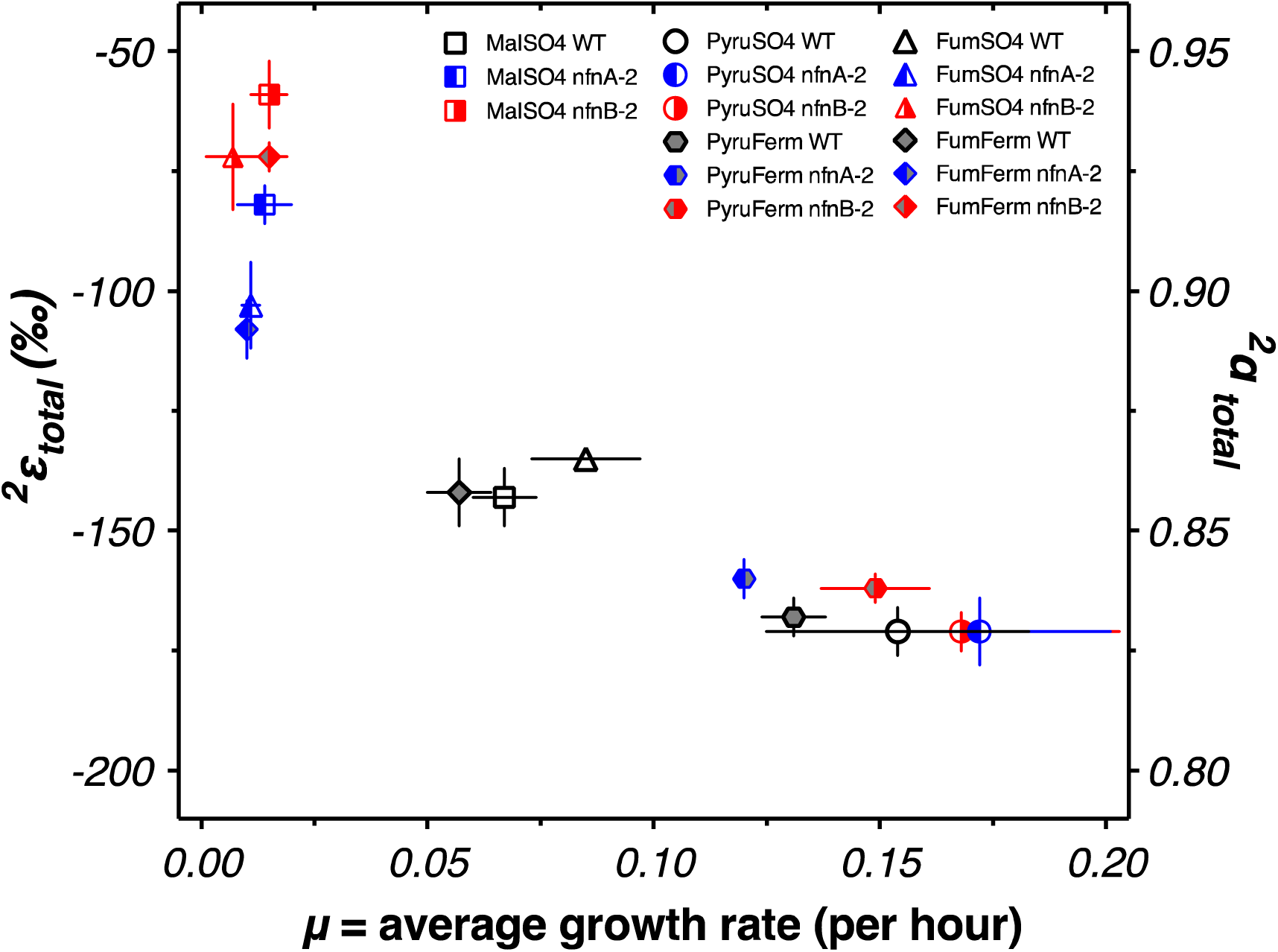
Average weighted growth rate (μ) versus the ^2^ε_total_ for all experiments.

## 4. Discussion

### 4.1 Hydrogen isotopes and intracellular electron flow

This study aims at improving our understanding of the influence of intracellular mechanisms that contribute to the ^2^H/^1^H ratios in lipids. Zhang et al. (2009) suggested that mechanisms related to the purine dinucleotide coenzymes NAD(P)H are central to determining δ^2^H_lipid_. In particular, that work suggested that H-isotopic fractionation by transhydrogenase was one potential mechanism for changing the H-isotopic composition of the transferable hydride on NAD(P)H. This is mainly due to the observations that NADPH directly provides up to 50% of lipid hydrogen (Robins et al., 2003; Saito et al., 1980; Schmidt et al., 2003), and predictions that δ^2^H_NADPH_ and abundance may vary with growth condition.

*Desulfovibrio* can produce NADPH via a number of mechanisms. Gram negative bacteria synthesize NADP^+^ from NAD^+^ by ATP-requiring NAD-kinase, but can not convert NADH to NADPH via this mechanism (Kawai and Murata, 2008). A number of metabolic reactions produce NADPH by reducing NADP^+^. Major mechanisms include enzymes within the tricarboxylic acid cycle, the oxidative pentose phosphate pathway, and mixed acid fermentation pathways. The relative importance of each of these mechanisms varies by substrate, and potential differences δ^2^H_NADPH_ produced by these mechanisms has been invoked as a major reason for differences in δ^2^H_lipid_ produced by organisms grown on various substrates (Zhang et al., 2009). Transhydrogenases are another mechanism for NADPH production. In addition to the Nfn family of transhydrogenase found in aerobes (Wang et al., 2010), two other families of transhydrogenases are common in aerobes: the proton-translocating transhydrogenase PntAB, and the energy-independent transhydrogenase UdhA (Sauer et al., 2004). These two enzymes have been discussed as a potential mechanism of influencing the δ^2^H of lipids (Dawson et al., 2015; Osburn et al., 2016; Zhang et al., 2009). Major mechanisms of NADPH production are summarized by Spaans et al. (2015), and those mechanisms relevant to *Desulfovibrio alaskensis* G20 are summarized in Table S2.

Previous work on hydrogen isotope fractionation in sulfate reducing bacteria (SRB) includes studies of pure cultures of *Desulfobacterium autotrophicum* (Campbell et al., 2009; Osburn, 2013), *Desulfobacter hydrogenophilus* and *Desulfovibrio alaskensis* G20 (Osburn, 2013) and of *Desulfococcus multivorans* in pure culture and in co-culture with a methanogen (Dawson et al., 2015). Results contrast with those obtained from aerobes, in which growth on different carbon sources results in a large range of ^2^ε_total_ (Zhang et al., 2009). *D. Autotrophicum* shows a smaller range in ^2^ε_total_ (~45‰) during heterotrophic growth on acetate, succinate, pyruvate, glucose, or formate, or autotrophic growth on H_2_/CO_2_, yet there are large differences in the ^2^ε_lipid_ of individual fatty acids (differences of >100 ‰) (Campbell et al., 2009; Osburn, 2013). *D. hydrogenophilus* and *D. multivorans* grown in pure cultures generate an ~80‰ range in ^2^ε_total_ during heterotrophic growth; though when *D. multivorans* was growing in co-culture with the methanogen *Methanosarcina acetivorans* it produced a more muted range in ^2^ε_total_ of ~36‰ (Dawson et al., 2015; Osburn, 2013).

In sulfate reducing bacteria, transhydrogenase NfnAB plays an important role in energy metabolism (Pereira et al., 2011; Price et al., 2014). If the magnitude of hydrogen isotope fractionation imparted by this transhydrogenase is large, similar to other transhydrogenases, then the activity of this enzyme may play a large role in setting _δ_^2^H_lipid_. This role may extend to the observed variations in δ^2^H_lipid_ as a function of substrate. In this model, NAD(P)H is produced or consumed by a variety of metabolic reactions in the cell (Table S2), but cycling of NAD(P)H through NfnAB could play a dominant role in determining the δ^2^H of NAD(P)H. The δ^2^H_lipid_ might be closely coupled to the size of the pools of oxidized and reduced purine dinucleotide coenzymes, rather than simply a function of changes in NAD(P)H δ^2^H. This could be a function of a Rayleigh distillation as NADPH is consumed. Alternatively, enzymes such as enoyl-ACP reductase, which are responsible for hydride transfer during fatty acid biosynthesis may be capable of utilizing either NADH and NADPH (Bergler et al., 1996), and the preferred substrate may change depending on their relative pool size. The perturbation of NfnAB in mutant strains would be expected to affect the relative sizes of these pools, and could help explain the observed patterns in lipids.

If NfnAB is important in determining δ^2^H_lipid_ in SRB, it could also be significant in other anaerobes as well. NfnAB genes are widely distributed in anaerobes, particularly in Deltaproteobacteria, Thermotogae, Clostridia, and methanogenic archaea (Buckel and Thauer, 2013), and thus far ubiquitous in sulfate-reducing bacteria (this study, Pereira et al., 2011). Figure 7 shows the relationship of NfnAB sequences from a range of anaerobes. Interestingly, all the sulfate reducing bacteria measure for lipid/water H-isotope fractionation to-date (Campbell et al., 2009; Dawson et al., 2015; Osburn et al., 2016; Sessions et al., 1999) contain some form of the NfnAB. While *nfnAB* sequences tend to cluster phylogenetically, the gene tree shown in Figure 7 identifies potential lateral gene transfer events among the SRB. The NfnAB genes from *Desulfobulbus propionicus* and *Syntrophobacter fumaroxidans* are more similar to those of the methanogenic archaea rather than the other SRB in the Deltaproteobacteria (including *D*. *alaskensis* G20), which cluster together. Similar to *D. alaskensis* G20, *D. propionicus* and *S*. *fumaroxidans* are SRB that are capable of fermentative growth on compounds such pyruvate (both) and fumarate (*S. fumaroxidans*).

**Figure 7.**
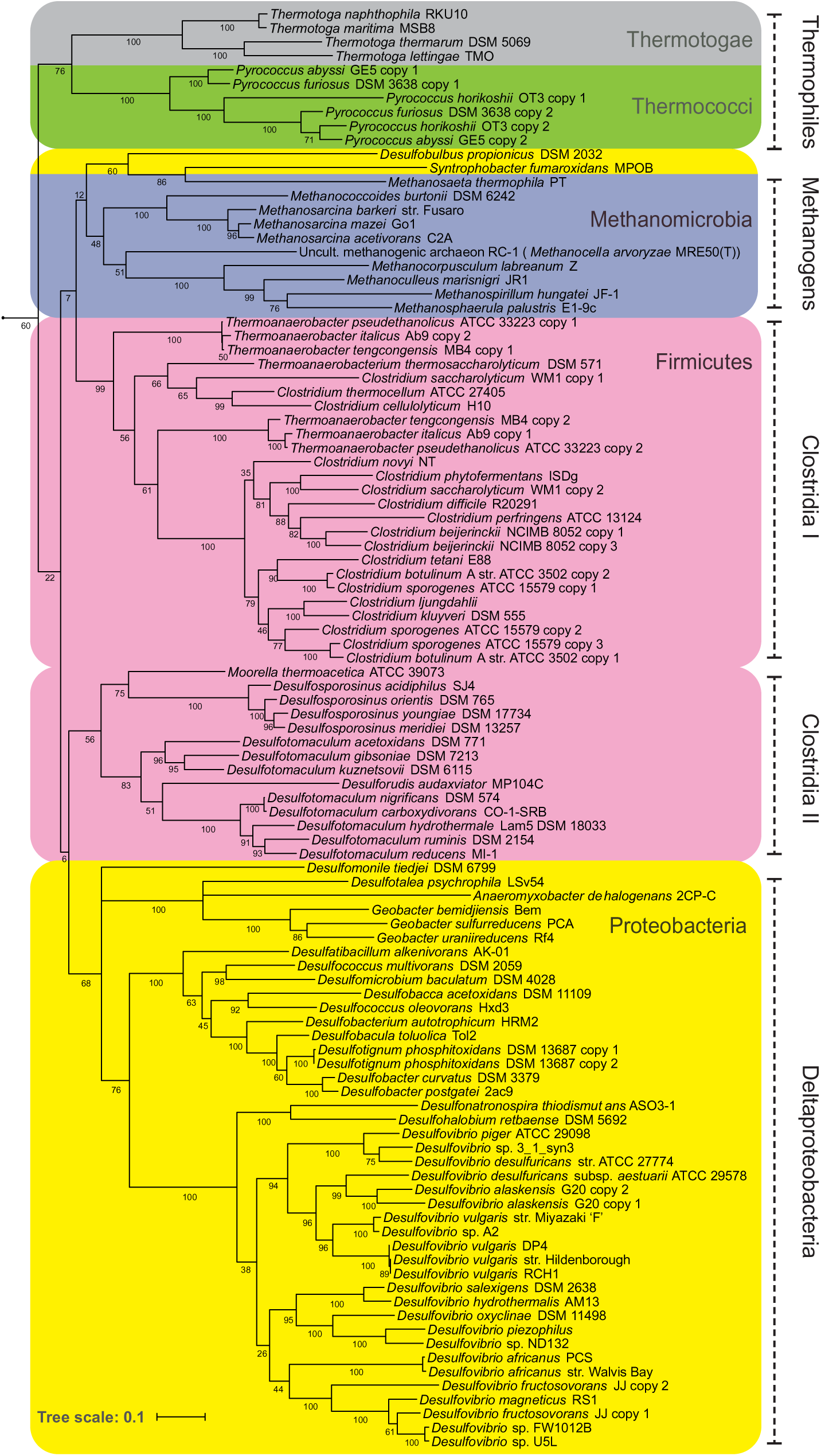
Maximum likelihood phylogenetic tree of NfnAB using amino acid sequences taken from the sequenced genomes of known anaerobes. Each branch is colored by Phylum. Bootstrap values (out of 100) are shown at each branch point.

In *D. alaskensis* G20, the catalytic role of the NfnAB transhydrogenase shows a relationship to the δ^2^H_lipid_ patterns. During growth on pyruvate/sulfate, electrons are predicted to flow from pyruvate to ferredoxin, then from ferredoxin through NfnAB to produce reduced NADPH (Price et al., 2014). That study pointed out that the reaction of transhydrogenase is probably required to produce sufficient NADPH for biosynthesis, even though the experiments were done in the presence of yeast extract, which minimized the importance of this reaction. In contrast, our isotopic experiments used a defined medium lacking yeast extract, so the importance of the transhydrogenase reaction would not be minimized. For both the wild type and mutant, growth on pyruvate/sulfate produced lipids that uniformly had the most negative ^2^ε_total_ across all our experiments. This experiment showed no phenotype for the *nfn* mutants, either in isotopic composition or in growth rate.

Sulfate reduction using malate likely employs NfnAB-2 to produce NADPH by oxidizing NADH and reduced ferredoxin (Price et al., 2014). Fumarate respiration operates in a manner similar to malate. These two substrates can be interconverted by fumarase (Price et al., 2014), so this similarity is likely to be related to similar growth and electron flow. Growth on each of these substrates produces similar patterns in hydrogen isotope fractionation. In each case, the wild type strain produces lipids with ^2^ε_total_ near −140‰, which is not as depleted in deuterium as lipids produced during growth on pyruvate. In contrast to growth on pyruvate, the mutant strains produce substantially less depleted lipids, resulting in a lesser degree of fractionation (^2^ε_total_) than the wild type. One explanation for this difference is that the mutation of one paralog of NfnAB in this strain changed the ratio of reduced to oxidized dinucleotides in the cell, with a higher ratio of NADPH to NADP^+^ and a lower ratio of NADH to NAD^+^. A second possibility is that the change in ^2^ε_total_ is a consequence of the growth defect of the mutant strains.

Previous work has investigated the relationship of growth rate to ^2^ε_total_. Zhang et al. (2009) did not observe a systematic relationship in the aerobic organisms that they studied. However, a negative relationship was observed between growth rate and the water-alkenone hydrogen isotope fractionation in the coccolithophores *Emiliania huxleyi* and *Gephyrocapsa oceanica* (Schouten et al., 2006). This observation is similar to that reported here, although the slope is steeper for *D. alaskensis* G20. Microbial lipids were recently reported to change their δ^2^H_lipid_ with growth phase (Heinzelmann et al., 2015), although this effect was relatively minor. Algal lipids have been reported to modulate _δ_^2^H_lipid_ as a function of physiological state (Estep and Hoering, 1980; Romero-Viana et al., 2013). Each of these relationships could be conceivably related to changes in the turnover rate or ratios of intracellular metabolites, but specific metabolomics data elucidating these relationships has yet to be produced.

Fermentation of pyruvate by *D. alaskensis* G20 likely involves the reduction of pyruvate with NADH by malic enzyme (ME; Dde_1253) to malate (Meyer et al., 2014), which is then dehydrated by fumarase to fumarate, and then reduced to succinate (*see* Price et al. 2014). The oxidative branch of this fermentation involves the transformation of pyruvate to acetate, which reduces ferredoxin. Reduced ferredoxin is recycled via flavin-based electron bifurcation, catalyzed by Hdr-Flox-1 (Meyer et al., 2014), but may also interact with NfnAB in the same way as during pyruvate respiration, in an electron confurcation reaction involving NADH, producing NADPH. Wild type *D. alaskensis* G20 grown by pyruvate fermentation produced lipids that showed ^2^ε_total_ comparable to that produced during pyruvate respiration. However, the *nfnAB-2* mutants grown by pyruvate fermentation generated an ^2^ε_total_ slightly more enriched than the wild type (Figure 4, mutants: −160‰ and −162‰, relative to WT: −168‰). If the *nfnAB-2* mutation inhibited NADPH formation, this pattern is opposite of that seen in other experiments, because it results in a less fractionation (a less negative ^2^ε_total_) at a lower predicted NADPH/NADP ratio. Nonetheless, the shift is minimal (<10‰) and the important reactions in ferredoxin recycling may be complicating this interpretation. Unlike the result in Meyer et al. (2014), *nfnAB-2* mutants in our experiments did not show a growth defect on pyruvate fermentation. This may in part be related to partial pressures of H_2_ produced by growing strains (not monitored herein). Alternatively, this may be due to the presence of the second transhydrogenase in G20, *nfnAB-1*, as highlighted above with respect to the pyruvate/sulfate experiments.

Fumarate fermentation in sulfate reducing bacteria is not well studied, but it is likely a complex metabolism. Wild type *D. alaskensis* G20 has nearly identical growth rates during fumarate fermentation and respiration. The *nfn* mutants grow more slowly than the wild type, but show little difference between fumarate fermentation and respiration. Respiration and fermentation of fumarate, along with respiration of malate, show nearly identical patterns in growth rate and in ^2^ε_total_ for each of our three strains (Figure 4). This suggests an underlying mechanism uniting these growth conditions.

Fatty acid profiles across the three strains and five conditions show some correlations with the total isotopic fractionation (Figure S3). The fractional abundance of branched chain lipids, particularly anteiso-C17:0, is positively correlated with the fractionation and negatively correlated with growth rate. We can reason two ways in which changing δ^2^H values of fatty acids may result in a less negative ^2^ε_total_ (Figure S2). The isotopic ordering of lipids may be altered, resulting in a net change in ^2^ε_total_; alternatively, there could be a consistent shift in the ^2^ε_lipid_ of most or all lipids. Data shown in Figures 3 and 5 rule out the first option, and show that almost all individual ^2^ε_lipid_ change in concert between conditions. This suggests that the driving mechanism for changes in 2ε relates to total process relevant to all lipids. Processes related to the production and consumption of NADPH are consistent with this role.

Integrating data from all five experimental conditions and three strains suggests a relationship between growth rate and ^2^ε_total_ (Figure 6). We do not yet have a theoretical prediction for the nature of this relationship, however the relationship is consistent with a linear, exponential decay, or hyperbolic relationship between growth rate and isotope fractionation. This pattern is similar, although opposite in sign, to that seen in sulfur isotope fractionation imposed by SRB during dissimilatory sulfate reduction (Chambers et al., 1975; Kaplan and Rittenberg, 1964; Leavitt et al., 2013; Sim et al., 2011, 2013).

Models aimed at addressing the growth rate—fractionation relationship in sulfur isotopes have focused on ratios of intracellular metabolites and redox state (Bradley et al., 2016; Wing and Halevy, 2014). Similar controls could be at work in controlling hydrogen isotope fractionation: ratios of NAD(P)H/NAD(P)^+^ and intracellular redox state are related and the partitioning of hydrogen between these pools could exert a direct effect on _δ_^2^H_lipid_. However, we cannot rule out the possibility that the correlation with growth rate is fortuitous, and correlations between δ^2^H_lipid_ and growth rate have not been observed in other studies (Dawson et al., 2015; Zhang et al., 2009). Ongoing work is aimed at testing this hypothesis, through growth of SRB in chemostats. If this hypothesis holds, then ^2^ε_total_ and δ^2^H_lipid_ of sulfate reducers may be able to provide a critical constraint on the interpretation of sulfur isotope patterns in natural systems, such as marine sediments and anoxic water columns.

Transhydrogenases related to *nfnAB* are widely distributed in anaerobes (Figure 7), and reactions catalyzed by this class of transhydrogenase may influence sedimentary lipid H-isotopic distributions in a wide range of natural settings. The metabolic role of NfnAB has been investigated in other anaerobes, notably thermophilic Clostridia (Lo et al., 2015), and similar studies of hydrogen isotope fractionation using these strains may indicate whether the patterns uncovered here in sulfate reducers are more generally applicable throughout obligate anaerobes, and the microbial domains of life in general.

The biggest differences in δ^2^H that we observe in these experiments are not between growth on various substrates or between wild type and mutant, but among individual fatty acids grown in a single culture. Understanding the biosynthetic mechanisms that are driving these differences will be key to the interpretations of the isotopic compositions of sedimentary fatty acids. The isotopic ordering is consistent, and differences in ^2^ε_total_ are not driven by changing abundances of lipids with extreme δ^2^H_lipid_, but by systematic changes across all lipids (Figure 5, S2).

The enrichment in δ^2^H_lipid_ of saturated fatty acids relative to their unsaturated homologues may be tied to biosynthesis. During fatty acid elongation, double bonds are introduced during each successive two-carbon addition. These *trans* double bonds are reduced by enoyl-ACP reductase, FabI (Kass and Bloch, 1967). A *trans* to *cis* configurational change can be introduced to the 10-carbon intermediate, preventing the function of FabI and conserving the double bond during further chain elongation. Fatty acids that have undergone this conversion are depleted in δ^2^H_lipid_, suggesting that the hydrogen transferred by FabI during fatty acid elongation is enriched in δ^2^H relative to average δ^2^H_lipid_. However, we cannot rule out the possibility that saturated and unsaturated fatty acids were produced at different times in the growth of our cultures and that the isotopic differences may reflect a different process than that articulated here.

Another observation necessitating explanation is the depletion in δ^2^H_lipid_ of anteiso-branched fatty acids relative to the straight chain fatty acids (Figure S2). Biosynthesis offers a potential explanation here as well. Straight-chain fatty acids are extended two carbons at a time by successive transfers of acetyl units (transferred as malonyl-ACP with the loss of CO_2_ during each transfer; ACP = acyl carrier protein). The primer for this chain extension is acetyl-CoA in the case of straight chain fatty acids, but differs for iso-and anteiso-branched fatty acids (Kaneda, 1991). Even-numbered iso-branched fatty acids use isobutyryl-CoA (derived from valine) as a primer, while odd-numbered iso-branched fatty acids use isovaleryl-CoA (derived from leucine). Even-numbered anteiso-branched fatty acids use 2-methylbutyryl-CoA (derived from isoleucine) as a primer (Kaneda, 1991). One possible explanation for a δ^2^H-depletion in anteiso-branched fatty acids is a depletion in the δ^2^H content of isoleucine. Compound-specific analysis of δ^2^H of amino acids has only recently been developed, but initial results suggest large differences in the δ^2^H content of various amino acids (Fogel et al., 2015). This new analytical technique could prove powerful in understanding the diversity of δ^2^H_lipid_ values produced by individual organisms.

## 5. Conclusions

The magnitude of hydrogen isotope fractionation in *D. alaskensis* G20 is influenced by the growth substrate, with growth on pyruvate exhibiting a different isotopic phenotype than growth on other substrates. Large differences are observed in the δ^2^H_lipid_ among individual lipids under all conditions. These differences may relate to biosynthesis, but are not fully accounted for. Wild type and *nfnAB-2* mutants show large differences in ^2^ε_total_ under conditions in which NfnAB-2 is predicted to play a significant role in energy conservation. This phenotype was observed across the entirety of the *D. alaskensis* G20 fatty acid profile. While ^2^ε_total_ correlates with modest changes in the fatty acids produced, it cannot be accounted for by changes in the abundance of individual lipids. These changes in apparent fractionation indicate a role for NfnAB-2 in determining both growth rate and δ^2^H_lipid_ for *D. alaskensis* G20, particularly when grown on malate or fumarate. NfnAB is widely distributed in anaerobes, and may play a role in determining δ^2^H_lipid_ in other organisms. Future work will aim to isolate these variables and further strengthen our understanding of the roles of growth and metabolic rate, substrate-induced differences in energy conservation pathways, and expression of transhydrogenase as key factors in the determining the hydrogen isotope ratios of lipids.

## 6. Acknowledgements

We thank Dr. A. Deutschbauer and Dr. J. Ray (Lawrence Berkeley National Laboratory) for providing *D. alaskensis* G20 mutant and wildtype strains, Dr. M. Osburn (Northwestern University) for water H isotope measurements and discussions of our data, and Dr. M. Lefticariu (Southern Illinois University, Carbondale) for external verification of lab H isotope standards. We also thank undergraduate researchers C. Wallace and L. Johnson (Washington University in St. Louis, WashU) for laboratory assistance. W. Leavitt acknowledges Washington University for the Steve J. Fossett Postdoctoral Fellowship. We thank Shuhei Ono for editorial support and two reviewers whose comments resulted in substantial improvements to the manuscript. T. Flynn acknowledges support from the Subsurface Science Scientific Focus Area at Argonne National Laboratory supported by the Subsurface Biogeochemical Research Program, U.S. Department of Energy (DOE) Office of Science, Office of Biological and Environmental Research, under DOE contract DE-AC02-06CH11357. This work was funded by NASA Exobiology grant 13-EXO13-0082.

---

**Figure S1.**
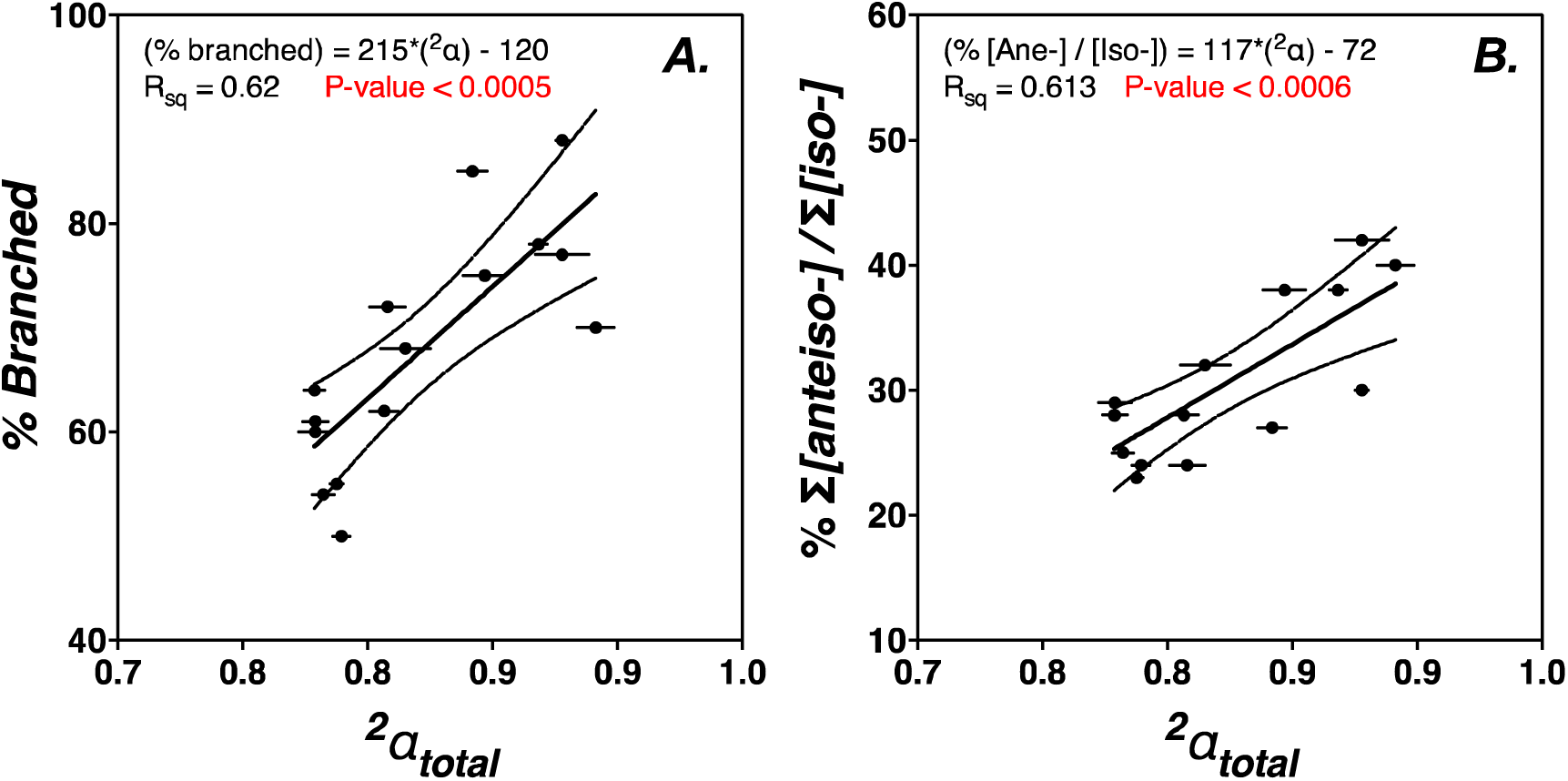
The mass-weighted fractionation between lipids and water versus the proportion of (A) branched fatty acids, (B) the ratio of anteiso‐ to iso‐ branched fatty acids.

**Figure S2.**
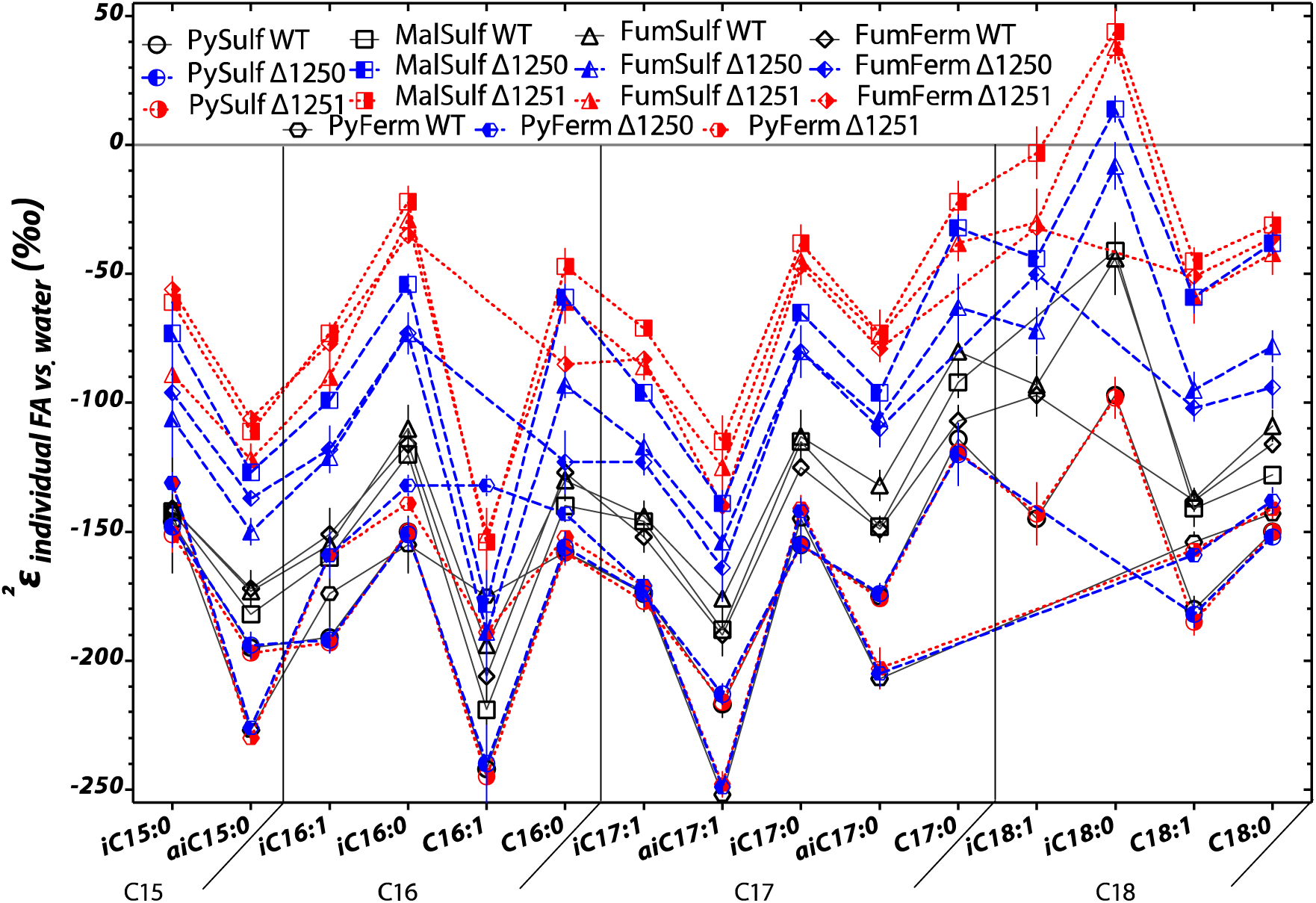
Hydrogen isotope values for individual fatty acids relative to medium water. These are the same values as in Figure 5, but here plotted by compound (x-axis) and legend-coded by experiment. Each symbol represents the mean of biological replicates (N = 2, and technical replication *n*_avg._ = 3, range 1 to 6), with SEM.

**Figure S3.**
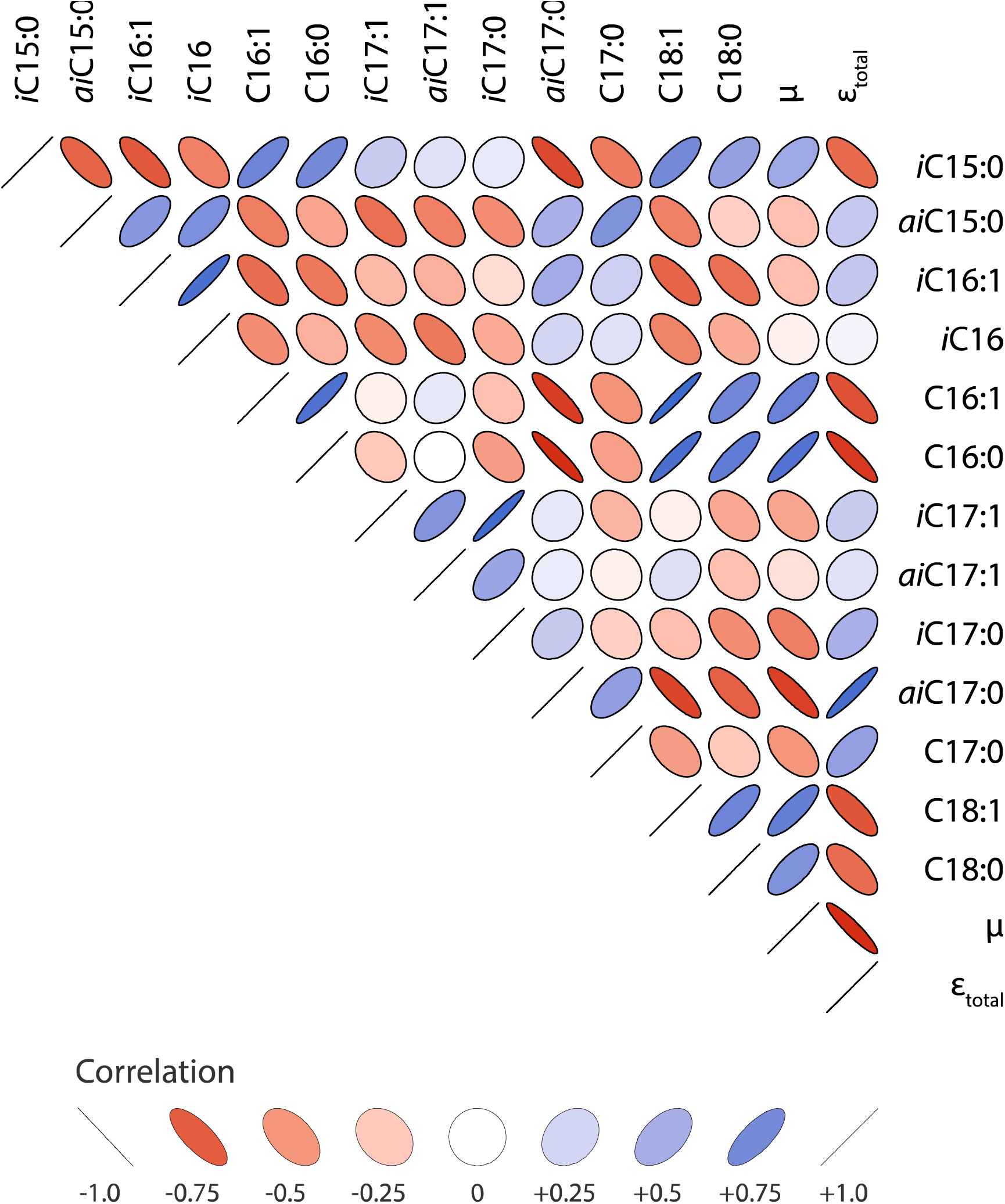
Representation of Pearson correlation indices for each pair of variables. Width of ellipses indicates the strength of the correlation, with narrow ellipses indicating a strong correlation and circles indicating no correlation. Darker blues are stronger positive correlations, darker reds are stronger negative correlations, with white indicating no correlation.

**Table S1.** FAME identifications based on mass spectra and retention times

| <u>RI</u> | <u>shorthand</u> | <u>M<sup>+</sup></u> | <u>Free fatty acid</u> |
| --- | --- | --- | --- |
| 1687 | i-C14:0 | 242 | 12-methyl tridecanoic acid |
| 1790 | i-C15:0 | 256 | 13-methyl tetradecanoic acid |
| 1798 | a-C15:0 | 256 | 12-methyl tetradecanoic acid |
| 1871 | i-C16:1 | 268 | 14-methyl pentadec-9-enoic acid |
| 1892 | i-C16:0 | 270 | 14-methyl pentadecanoic acid |
| 1905 | C16:1 | 268 | hexadec-9-enoic acid |
| 1928 | C16:0 | 270 | hexadecanoic acid |
| 1967 | i-C17:1 | 282 | 15-methyl hexadec-9-enoic acid |
| 1976 | a-C17:1 | 282 | 14-methyl hexadec-9-enoic acid |
| 1991 | i-C17:0 | 284 | 15-methyl hexadecanoic acid |
| 2000 | a-C17:0 | 284 | 14-methyl hexadecanoic acid |
| 2004 | C17:1 | 282 | heptadecenoic acid |
| 2010 | C17:1 | 282 | heptadecenoic acid |
| 2027 | C17:0 | 284 | heptadecanoic acid |
| 2070 | i-C18:1 | 296 | 16-methyl heptadec-11-enoic acid |
| 2090 | i-C18:0 | 298 | 16-methyl heptadecanoic acid |
| 2108 | C18:1 | 296 | octadec-11-enoic acid |
| 2132 | C18:0 | 298 | octadecanoic acid |
| 2738 | C24 | 382 | tetracosanoic acid (standard) |

**Table S2.** Major mechanisms of NADPH production relevant to *D. alaskensis* G20

| <u>Enzyme</u> | <u>EC number</u> | <u>G20 locus</u> | <u>name</u> |
| --- | --- | --- | --- |
| G6PDH | 1.1.1.49 | Dde_3471 | glucose-6-phosphate 1-dehydrogenase |
| 6PGDH | 1.1.1.44 | Dde_3470 | 6-phosphogluconate dehydrogenase |
| IDH | 1.1.1.42 | Dde_3476 | Isocitrate dehydrogenase |
| ME | 1.1.1.40 | Dde_1253 | malic enzyme |
|  |  |  | non-phosphorylating glyceraldehyde 3- |
| GAPN | 1.2.1.9 | nd | phosphate dehydrogenase |
| NADP+- |  | Dde_2342, | Glyceraldehyde-3-phosphate dehydrogenase, |
| GAPDH | 1.2.1.13 | Dde_3736 | type I |
|  | 1.1.1.47, |  |  |
| GDHs | 1.1.1.119 | nd | glucose dehydrogenase |
|  |  |  | energy-independent soluble |
| STH | 1.6.1.1 | nd | transhydrogenase |
|  |  |  | energy-dependent or proton- |
|  |  |  | translocating, membrane-bound |
| H+-TH | 1.6.1.2 | nd | transhydrogenase |
|  |  | Dde_3636, |  |
| FNR | 1.18.1.2 | Dde_1251 | ferredoxin-NADP reductase |
| SH | 1.12.1.3 | Dde_1212 | soluble hydrogenase |
| NADK | 2.7.1.23 | Dde_2618 | ATP-NAD/AcoX kinase |
| PDR | 1.8.1.8 | Dde_1301 | Protein-disulfide reductase |
|  |  | Dde_3635, |  |
| NfnAB1 | 1.6.1.4 | Dde_3636 | electron-bifurcating transhydrogenase |
|  |  | Dde_1250, |  |
| NfnAB2 | 1.6.1.4 | Dde1251 | electron-bifurcating transhydrogenase |

\*most of these reactions are derived from Table 1 in Spaans et al., 2015

